# Intradermal and intranasal vaccination with microparticulated ASFV proteins induces distinct immune profiles in pigs

**DOI:** 10.64898/2026.11.23.768517

**Authors:** Elena Coatu, Sonia Pina Pedrero, Aida Tort-Miró, Enrique Ezcurra, Paula Monleón, Marta Muñoz, María J Navas, Judith González-Oliver, Àlex Cobos, Miquel Nofrarias, Hèctor López-Laguna, Antonio Villaverde, Esther Vázquez, Anna Arís, Elena Garcia-Fruitós, Fernando Rodríguez, Francesc Accensi, Jordi Argilaguet, Sergio Montaner-Tarbes

**Author notes:** these authors equally contributed to this work (Jordi Argilaguet, Sergio Montaner-Tarbes).

## Abstract

African Swine Fever (ASF) is a major threat to the swine industry due to its severe socioeconomic impact and the lack of safe and effective vaccines. Although live attenuated vaccines (LAVs) can induce robust immunity, biosafety concerns restrict their use, making subunit vaccines the preferred alternative. In this context, understanding how vaccination routes modulate immunity is critical. A previous study using a viral vector-based vaccine, encoding for the ASFV antigens p30, p54, p72, p72c and CD2v, induced protection when combining intranasal (IN) and parenteral administrations. However, the relative contribution of each administration route to vaccine-induced immunity remains unclear. Here we investigated how independent and combined vaccination routes influence the immune responses induced by the same ASFV protein cocktail delivered as nanoparticle-based secretory granules. Following a prime-boost regime, pigs vaccinated via intradermal (ID) route seroconverted and developed strong systemic ASFV-specific IgG responses, despite inducing only modest ASFV-specific cellular responses. In contrast, IN administration alone failed to elicit detectable systemic humoral or cellular responses and did not improve the vaccine-induced immunogenicity when combined with the ID route. Following a challenge with a virulent ASFV strain, no protection was achieved in any of the immunized groups. Interestingly, intranasally vaccinated pigs exhibited an accelerated post-challenge ASFV-specific antibody boost in serum compared to controls, suggesting that vaccination through this mucosal route induced immune priming despite the absence of detectable systemic humoral responses before challenge. Overall, these findings reveal distinct route-dependent immune profiles elicited by the same ASFV antigen cocktail and highlight again that robust humoral responses alone are insufficient for protection, further supporting the importance of cellular immunity in vaccine-induced protection.

**Highlights:**

- Intradermal vaccination with the ASFV antigens p30, p54, p72, p72c and CD2v induced strong systemic antibody responses.
- Intranasal vaccination induced evidence of immune priming despite the absence of detectable systemic responses before challenge.
- Neither vaccination route conferred protection against virulent ASFV challenge.
- Robust antigen-specific antibody responses alone were insufficient for protection against ASFV.

## 1. Introduction

The development of subunit vaccines against African swine fever virus (ASFV) has advanced considerably in recent years, with several studies demonstrating promising degrees of protection using diverse approaches and antigen combinations [1], [2], [3], [4], [5]. However, vaccine-induced efficacies remain inconsistent and generally insufficient for field application, highlighting the need to optimize vaccination strategies. While the identification of protective antigens and correlates of protection remains a major challenge [6], increasing evidence suggests that vaccine efficacy also depends on factors such as antigen delivery systems and administration routes, which can significantly influence the quality, magnitude and tissue distribution of immune responses. In this context, a recent study using a combination of five adenovirus- vectored ASFV proteins (p30, p54, p72, p72c and CD2v) demonstrated solid protection against a natural ASFV challenge in farm pigs, supporting the inclusion of these antigens in vaccine formulations [4]. Interestingly, protection was achieved following simultaneous intramuscular and intranasal vaccine administration, although the relative contribution of each route was not investigated.

Administration route significantly influences the type of vaccine-induced immune responses, thereby affecting vaccine efficacy. Intramuscular (IM) inoculation has traditionally been the most widely used approach due to its practicality, despite muscle tissue being relatively poor in professional antigen-presenting cells [7]. In contrast, intradermal (ID) immunization often induces enhanced immune responses thanks to the high density of dendritic cells and efficient lymphatic access [8], [9], [10], [11]. Importantly, the efficacy of ID vaccination has also been demonstrated in several swine models [12], [13], [14]. However, parenteral vaccination strategies generally induce limited mucosal immunity, which may be particularly relevant against pathogens that can establish infection through mucosal surfaces, including ASFV [15], [16]. Consequently, intranasal (IN) immunization is emerging as a promising strategy to improve vaccine efficacy, as demonstrated both in murine models [17] and humans [18]. In pigs, IN vaccination has been shown to induce both mucosal humoral and cellular immune responses, contributing to the control of respiratory infections [19], [20], [21], [22]. However, the relative performance of intranasal and parenteral vaccination strategies is highly dependent on vaccine formulation and delivery platform [23], [24], [25], [26]. Despite the growing interest in mucosal vaccination, direct comparisons of administration routes in ASF vaccine research remain scarce and have been largely restricted to live attenuated vaccine strains [27], [28], [29]. Consequently, how different administration routes influence the immunogenicity of ASFV subunit vaccines remains largely unknown. Importantly, studies using naturally attenuated ASFV strains have shown that IN inoculation can confer higher levels of protection against virulent challenge than IM inoculation [30], [31], suggesting a potentially important role of mucosal immunity in protection against ASFV.

In this study we aimed to investigate how the vaccination route of an experimental ASFV subunit vaccine influences the immune responses and protection induced. To address this question, we selected the same five ASFV antigens (p30, p54, p72, p72c and CD2v) previously evaluated by Liu et al. in an adenovirus-vectored vaccine that conferred a robust protection following natural ASFV exposure [4]. However, similar antigen combinations had not previously been associated with such levels of protection in alternative vaccine platforms, thus requiring further validation of their protective potential in an independent formulation. Therefore, we formulated a subunit vaccine using nanoparticle-based secretory granules, a delivery platform with inherent immunostimulatory properties that has been previously shown to induce antigen- specific humoral and cellular immune responses following both IN and parenteral administration in mice and pigs [32], [33]. Using this platform, we evaluated the influence of ID, IN, and combined ID-IN vaccination on the immunogenicity and protective efficacy of this ASFV antigen combination in pigs. The results show that, in this model, ID administration induced stronger systemic immune responses than IN vaccination, although neither strategy conferred protection against virulent ASFV challenge. These findings further support the notion that, beyond the antigen composition, the quality of the immune response induced by different vaccination strategies is a critical determinant of protection against ASFV.

## 2. Methods

### 2.1. Ethics Statement

Animal care as well as the respective procedures were subjected to the Guidelines of the Good Experimental Practice and approved by the Ethics Committee on Animal Experimentation of the Generalitat de Catalunya. The project code for this experiment is CEA-OH/12433/1.

### 2.2. Plasmid Design for ASFV proteins

The plasmid constructs for the recombinant proteins p72, CD2v, p30 and p54 encoded by genes *pB646L, EP402R, CP204L* and *E183L* respectively, were designed based on the Georgia2007/1 strain (NCBI Reference Sequence: NC_044959.2). Protein p72c, encoded by *pB602L*, was obtained from *in-house* Georgia2007/1 sequence (GenBank: CAD2068442.1).

All proteins were modified to add a poly-His Tag (H6) at the C-terminal end. Additionally, p30 included a RK motif at the N-terminal end and CD2v protein was designed to have the transmembrane region replaced by a 15 amino acid linker sequence (G_4_S)_3_.

Plasmids RK4-p30-H6 and p54-H6 were provided by Gene Art (Thermo Scientific Fisher, Waltham, MA, USA) and subcloned in pET22b (Amp^R^) and pASK33-IBAPLUS (Amp^R^) (Novagen, Madison, WI, USA) expression vectors, respectively. Plasmid p72c- H6 was also subcloned in the pASK33-IBAPLUS expression vector.

The proteins p72-H6 and CD2v-H6 were ordered from GenScript (Piscataway, NJ, USA) based on the Georgia2007/1 sequence (GenBank: CAD2068442.1).

### 2.3. Protein synthesis in prokaryotic system

The expression vector was transformed by heat shock in competent *Escherichia coli* BL21 (DE3) strain (Thermo Scientific, EC0114) for p30 and p54, and in *E. coli* Rosetta (DE3) strain for p72c obtention. The transformed bacteria were grown in Luria-Bertani (LB) broth with carbenicillin at 50 ug/mL in 5 L flasks with 2.5 L of media at 37°C and 250rpm. When producing p72c protein, chloramphenicol at 34 ug/mL was added to maintain plasmid retention and selection stringency. The induction of the recombinant proteins began when the culture OD600 reached 0.5. For p30 protein induction was performed at 20°C and 200 rpm for 3 h with 0.1 mM isopropyl-β-d- thiogalactopyranoside (IPTG, Invitrogen, 15529-019). For p54 and p72c proteins, induction was done with 0.2 mg/ml anhydrotetracycline tetrachloride (AHT, Thermo Scientific, J66688.MB) at 20°C and 200 rpm, for 3 h when producing p72c and overnight when producing p54.

After protein induction, cells were harvested by centrifugation at 3,220 × g for 15 to 60 minutes and stored at -80°C until further use. Before purification, bacterial pellet were resuspended in ÄKTA Binding Buffer (20 mM Tris, 500 mM NaCl and 20 mM Imidazole) and disrupted at 20 kPsi with Constant Systems CF1 disruptor. PMSF (Phenylmethylsulfonyl fluoride, MERCK, 52332-5G) protease inhibitor at 1 mM was added to the lysate, and it was centrifuged at 15,000 × g for 45 minutes at 4°C. Later, DNase I (Roche, 4716728001) was added to the supernatant before filtering it through a 0.22 um PVDF membrane (Millipore, GVWP02500). Purification was performed by Immobilised Metal Affinity Chromatography (IMAC) in ÄKTA Start™ chromatography system (Cytiva) using 5 mL HisTrap High Performance columns (Cytiva, GE17-5248-02). The eluted proteins in ÄKTA Elution Buffer (20 mM Tris, 500 mM NaCl and 500 mM Imidazole) were dialyzed twice against a carbonate-bicarbonate buffer (166 mM NaCO_3_H, 333 mM NaCl). Finally, the dialyzed proteins were concentrated using 3000 MWCO Amicon Ultra-15 Centrifugal Filter Devices (Millipore, UFC900324) and the concentrated proteins were stored at -80°C until characterization and use.

### 2.4. Generation of nanoparticle-based secretory granules

The ASFV proteins p30, p54, p72 and p72c were successfully formulated as nanoparticle-based secretory granules following the methodology previously described by Bosch-Camós et al. [33]. In contrast, the CD2v was included in the vaccine formulation as a soluble recombinant protein, since its glycosylated structure was incompatible with the His-tag-driven assembly process required for secretory granule formation. Proteins p30, p54 and p72c were microparticulated in 150 ug aliquots by the addition of 10 mM ZnCl_2_ (Sigma Aldrich, 208086-100G), whereas the p72 required 25 mM ZnCl_2_ for efficient granule formation. Following a 10-minute incubation at room temperature, samples were centrifuged at 10,000 × g for 10 minutes. The resulting granules were stored at -80°C until use.

### 2.5. Western Blot

Proteins were characterized by western blot and SDS-PAGE electrophoresis. ASFV Proteins and ladders (BenchMark™ His-tagged Protein Standard - Thermo Scientific, LC5606 / PageRuler Prestained Protein Ladder - Thermo Scientific, 26616) were loaded in the NuPAGE^TM^ 10% Bis-Tris Gel (Thermo Scientific, NP0303BOX) and run for 45 minutes at 180 V and 250 mA in the XCell SureLock^TM^ Mini-Cell. After SDS- PAGE electrophoresis, membrane transfer was performed using the Mini Trans-Blot Electrophoretic Transfer Cell (BioRad, 1703930) for 2 h at 50V and 250 mA with the nitrocellulose membrane Amersham^TM^ Protran® Premium 0.45 um NC (Sigma-Aldrich, 10600003). As quality control for protein transfer, the membrane was stained with ATX Ponceau S red staining solution (Sigma-Aldrich, 09189). After overnight blocking at 4°C on an orbital shaker, the membrane was incubated with 1:100000 dilution of HRP- 6X His tag monoclonal antibody (Fine Test, FNSA-0019) for 1 hour. Finally, SuperSignal™ West Femto Maximum Sensitivity Substrate (Thermo Scientific, 34096) was used for specific His-tagged protein detection and the image was acquired on Fluorchem HD2, Alpha Innotech.

### 2.6. Virus

The highly virulent Georgia 2007/1 virus (genotype II) was kindly provided by Dr. Linda Dixon at The Pirbright Institute, UK, from the World Organization for Animal Health (WOAH) reference laboratory. The strain was produced in porcine alveolar macrophages (PAMs) procured by bronchoalveolar lavages from healthy pigs which tested negative by porcine circovirus type 2, porcine reproductive and respiratory syndrome virus (PRRS), and *Mycoplasma spp,* as previously described [34]. The virus was titrated by hemadsorption assay [35] and its titre was calculated as 50% hemagglutination activity units (HAU50)/mL, according to the Reed and Müench method [36].

### 2.7. Animal Experiment

The animal experiment was performed with twenty-four 5-week-old healthy Landrace x Large White piglets *(Sus scrofa domesticus)*, male and female, from Grup Batallé (Catalunya, Spain). Upon arrival at the biosafety level 3 (BSL-3) facility, the animals were randomly allocated to four groups of six piglets each. The pigs were then individually numbered and allowed to acclimate for one week before the start of the vaccination trial. Throughout the whole study pigs had food and water *ad libitum and*, to promote their natural foraging conduct, ropes and rubber floor toys were offered. Animals were observed daily and assessed for clinical signs, body condition, behaviour, and temperature. Weekly sampling included blood collection, nasal swabs and salivettes. The protein cocktail was inoculated following a primer-boost vaccination schedule at 0 and 14 dpv. Each vaccination dose contained 150 ug of each of the following microparticulated proteins: p30, p54, p72, p72c, except for CD2v that was in its soluble form.

Groups were organized according to the administration route. The control group (pigs 7 to 12) received sterile PBS (Corning, 21-040-CV) intranasally (2 mL; 1 mL per nostril) using a nasal atomizer and intradermally (500 µL) in the neck using a 20-G needle. Pigs 1 to 6 received the microparticulated protein cocktail by both routes using the same procedures. Pigs 13 to 18 and 19 to 24 received the vaccine candidate exclusively by the intranasal or intradermal routes, respectively, following the abovementioned procedure.

At 28 dpv animals were challenged with 10^5^ HAU of the highly virulent genotype II strain Georgia 2007/1 per pig. The inoculum was prepared in 2 ml of sterile PBS and administered intranasally using a 2 ml syringe with a nasal atomizer, dispensing 1 ml in each nostril. After the challenge, animals were euthanized when they reached the humane endpoint according to the animal care facilities using intravenous pentobarbital (140 mg/kg). Necropsies were performed to confirm the presence of ASF-compatible pathological lesions and to collect tissues (tracheobronchial and gastrohepatic lymph nodes). The scoring of the ASFV macroscopic lesions was performed according to Galindo-Cardiel et al. method [37].

### 2.8. Quantitative PCR for the Detection of ASFV

Viremia was quantified by qRT-PCR from blood and nasal swabs. The samples were processed to extract the viral DNA with the IndiMag Pathogen Kit (INIDICAL BIOSCIENCE, SP947457) following their own standard operational procedure. Isolated DNA was used as a template for qRT-PCR. Primers and probes targeting the ASFV PK gene were used [38] in combination with Path-ID qPCR Master Mix (Thermo Scientific, 4388644) and the 7500 Fast Real-Time PCR System. Samples were considered positive if they were above the quantification limit (LOQ > 6 GEC/uL). A set of quantified serial dilutions containing the PK gene was used as standard and positive and negative controls were included to validate the DNA extraction and qRT- PCR.

### 2.9. Enzyme-Linked Immunosorbent Assay (ELISA)

ASFV-specific antibodies in pig serum were measured by the WOAH-approved ELISA test following their own standard operational procedure [39]. For each individual protein produced, an in-house ELISA test was standardized. For both, flat bottom high binding 96-well plates were coated (Corning, 3590) with either AgL1 (1:1600 dilution in carbonate-bicarbonate coating buffer) or ASFV proteins (p54, p30, p72, p72c and CD2v) at 0.5 ug/mL. Only for the protein antigenicity ELISA, protein coating was done with 20 ug/mL. Plates were blocked with PBS 1X - 5% non-fat dry milk and then washed with PBS 1X 0.05% Tween 20. ASFV-protein specific antibodies (against p54, p30, p72, p72c and CD2v) were detected using a serum dilution of 1:100 in blocking buffer. All samples and controls were run in technical duplicates. To calculate the endpoint-titer for the ASFV virus-specific and ASFV-individual protein antibodies, samples were serially diluted from 1/100 to 1/1.562.500 and 1/50 to 1/6400 respectively.

Positive sera or nasal swabs were detected using the following secondary peroxidase- conjugate antibodies: rabbit anti-pig IgG at 1:20000 dilution (Sigma-Aldrich, A5670), goat anti-pig IgM at 1:100000 (BioRad, AAI48P) or goat anti-pig IgA at 1:10000 (Bethyl, A100-102A). As substrate for the peroxidase reaction, soluble 3,3’,5,5’- tetramethylbenzidine (TMB, Sigma-Aldrich, T4444) was used. To stop the peroxidase reaction, H2SO4 at 1 N was used. Finally, plates were read at 450 nm. Samples ODs were normalized against the positive control, so results are displayed as ELISA Index. The cut-off value or threshold was defined for each ELISA as the mean of samples replicates at 0dpv plus three times the standard deviation of the negative control. This value is depicted as a dashed grey line in the ELISA plots.

### 2.10. IFN-ɣ *enzyme-linked immunosorbent spot (ELISpot) assay*

The IFN-ɣ-ELISpot assay was performed as previously described [40] in 96-well Multiscreen^®^ nitrocellulose plates (Millipore, MAIPS4510). First, ELISPOT membranes were activated by adding 50 uL of 35% ethanol and incubated 1 minute at room temperature. All wells were washed with sterile PBS 1X and coated with the capture antibody (purified mouse anti-pig IFN-ɣ clone P2G10, BD Pharmingen, 559961). PBMCs were isolated for ELISPOT and flow cytometry from porcine whole blood in EDTA tubes following a Histopaque® gradient (Sigma-Aldrich, 10771-100ML). After centrifugation, cell viability was assessed by trypan blue live staining using a Neubauer chamber. Cells were seeded in complete RPMI at 5×10^5^ cells/well and stimulated with either Georgia2007/1 at a MOI of 0.2 or the protein cocktail at a final concentration of 10 ug/mL and incubated for 16 hours at 37°C, 5% CO2. For negative and positive controls, RPMI media (Corning, 15-040-CV) and phytohemagglutinin in M form (PHA- M, Sigma-Aldrich, 11082132001, 5ug/mL) were used respectively.

Finally, IFN-ɣ was detected with a biotin mouse anti-porcine IFN-ɣ antibody (clone P2C11, BD Pharmingen, 559958) and Streptavidin-ALP (MABTECH, 3310-10-1000). Plates were read and counted with AID iSPOT ELISpot FLUOROSpot Reader (Advanced Imaging Devices GmbH, ELR08IFL) using the AID ELISpot Reader Software Version 7.0.

### 2.11. Flow Cytometry analysis

PBMCs were isolated as described above. For each animal and stimuli, 1×10^6^ fresh PBMCs were seeded per well in a round bottom 96-well plate (Corning, 3799). To assess perforin positive cells within T cell populations, cells were stimulated separately with Georgia 2007/01 at MOI 0.2 for 36 h at 37°C, with the protein cocktail or each protein separately at a final concentration of 10 ug/mL. For negative and positive control, complete RPMI media (Corning, 15-040-CV) and phorbol myristate acetate (PMA, 5 ng/mL) plus ionomycin (500 ng/mL) were used.

After 36 h stimulation, PBMCs were stained with Live/Dead® Fixable Dead Cell Stain Kit (Thermo Scientific, L34975) and washed two times with 150 uL of PBS 1X / 2% FBS. Each wash, samples were centrifugated at 350 × g for 3 minutes. Blocking of Fc receptors was done with 100 uL of PBS 1X / 5% porcine serum and incubated for 15 minutes at 4°C. Samples were centrifuged at 400 × g for 3 minutes and washed twice with 150 uL PBS 1X / 2% FBS. After, the extracellular staining was performed with 50 uL of the antibody mix containing mouse anti-pig CD3ε (PE-Cy7, Clone BB23-8E6- 8C8, BD Biosciences, 561477), mouse anti-pig CD4α (PercP-Cy5.5, Clone 74-12-4, BD Biosciences, 561474), rat anti-pig γδ T Lymphocytes (APC, Clone MAC320, BD Biosciences, 561482), anti-pig CD8α (FITC, Clone 76-2-11, BD Biosciences, 551303) and mouse anti-pig CD8β chain (R-PE, Clone PPT23, Bio-Rad Laboratories, MCA5954PE). This incubation was carried out on ice for 20 min in the absence of light.

Cells were washed with 150 uL of PBS 1X / 2% FBS and centrifuged at 400 × g for 3 minutes. Fixation and permeabilization step was done with BD Cytofix/Cytoperm Kit (BD Biosciences, 554714) as instructed by the supplier and cells were washed with 150 uL of BD Perm/Wash^TM^ 1X by centrifugated at 400 × g for 3 minutes. Finally, cells were stained intracellularly with mouse anti-human/pig perforin (eFluor^TM^ 450, clone delta G9, eBioscience, 48-9994-42) and incubated on ice for 30 min in the absence of light. Following a wash with 150 uL of BD Perm/Wash^TM^ 1X and centrifugation at 400 × g for 3 minutes, cells were resuspended in PBS 1X / 2% FBS. Acquisition was performed on a BD FACSDiscover™ S8 Cell Sorter with BD CellView™ Image Technology and BD SpectralFX™ Technology (BD Life Sciences). Flow cytometry acquired data was analysed with FlowJo Version 10.10.0 (FlowJo LLC).

### 2.12. Statistical analysis

Statistical analyses were performed with GraphPad Prism Version 9.0.2. Normality of data distribution was tested by the Shapiro-Wilk test. Each statistic test used to further analyse data is mentioned in the corresponding figure legend as well as its p-values. Briefly, for paired t-tests statistical difference was set up at: *p value < 0.05; **p value < 0.005 and ***p value < 0.0005. For one-way and two-way ANOVA the statistical difference was set up at: *p value < 0.05; **p value < 0.01; ***p value < 0.001 and ****p value < 0.0001.

## 3. Results

### 3.1. Robust systemic humoral response following intradermal administration of the microparticulated antigen cocktail

We previously demonstrated that immunization with nanoparticle-based secretory granules is effective at inducing ASFV-specific immune responses in pigs (Bosch- Camós et al., 2024). Here, we evaluated how ID and IN vaccination routes influence the induced ASFV-specific immune responses. With this aim, the ASFV proteins p30, p54, p72, p72c and CD2v were selected as vaccine antigens, and the preservation of antigenic epitopes was confirmed by ELISA using ASFV-positive sera (Fig. S2). Pigs were vaccinated with the protein cocktail in a prime-boost regimen (Fig. 1A), and systemic humoral and cellular responses were analysed. No adverse effects were observed following vaccination regardless of the administration route. ASFV-specific IgG antibodies in serum were detected only in the ID and ID+IN groups, becoming significant after the booster immunization (Fig. 1B) and reaching the highest endpoint titres at 28 dpv (Fig. 1C), whereas IN vaccination alone failed to induce detectable systemic responses (Fig. 1B). Protein-specific ELISAs confirmed that antibody responses were directed against all antigens included in the vaccine cocktail, although marked differences in immunogenicity were observed (Fig. 2). The strongest responses were detected against p30, p54, and p72c, whereas antibody recognition of p72 and CD2v were comparatively lower. Consistent with the ASFV-specific ELISA, antigen-specific responses were detected only in the ID and ID+IN groups. Finally, no ASFV-specific IgM responses were detected in serum throughout the study period (Fig. S3).

**Fig. 1.**
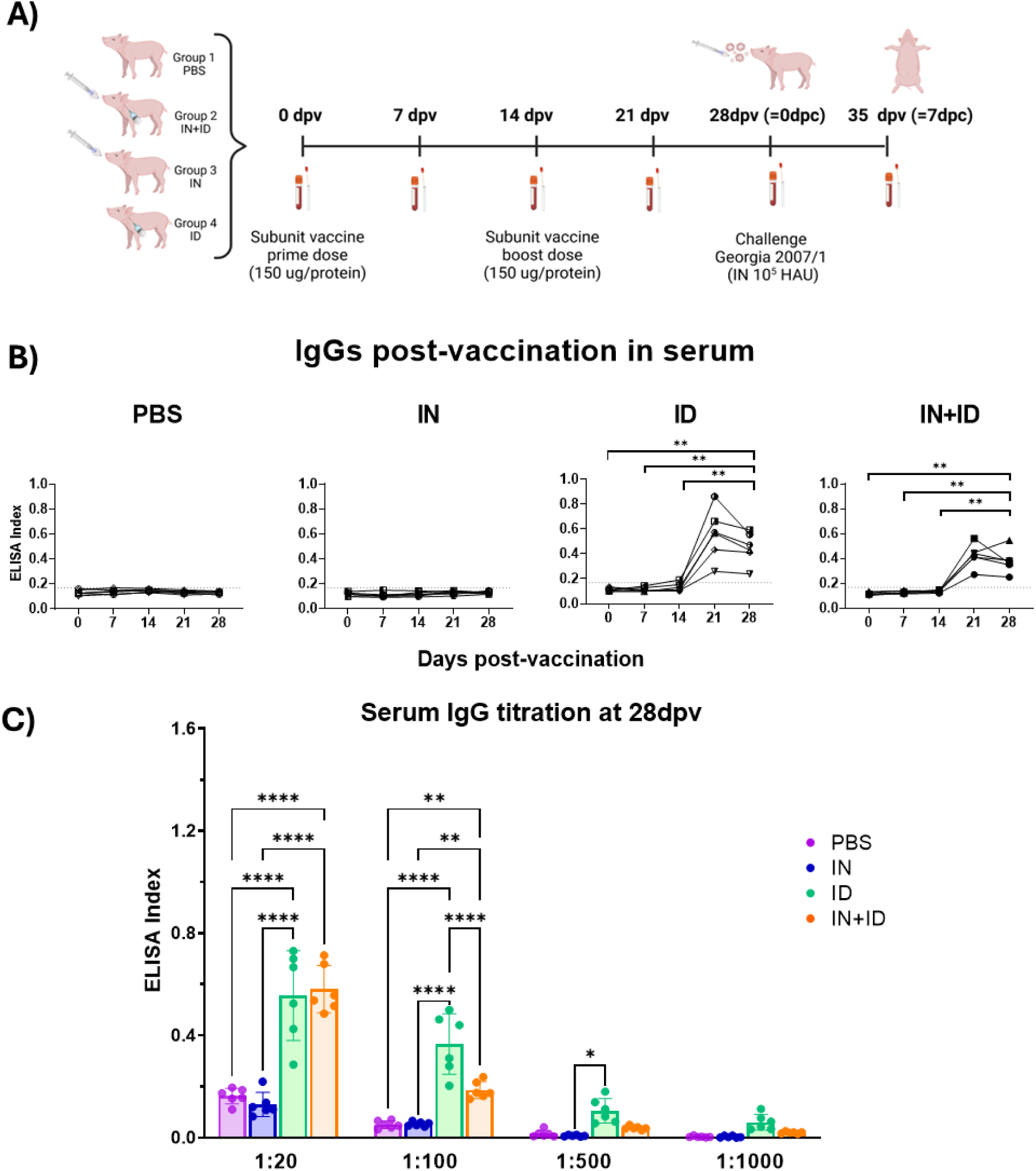
Intradermal vaccination induces robust ASFV-specific IgG antibody responses. (A) Experimental design. Four groups of piglets (n=6 per group) were immunized following a prime-boost regimen (days 0 and 14) with a vaccine cocktail containing microparticulated p30, p54, p72 and p72c, and soluble CD2v. Animals were vaccinated by the intradermal (ID), intranasal (IN) or combined (ID+IN) routes. Control animals were inoculated with PBS. Created with BioRender. (B) ASF-specific IgGs antibodies in serum from 0 to 28 dpv by ELISA. (C) ASFV-specific serum IgG responses at 28 dpv across serial serum dilutions. Statistical significance was determined by a one-way (B) or two-way (C) ANOVA (\**P* < 0.05, \*\**P* < 0.01, \*\*\**P* < 0.001 and \*\*\*\**P* < 0.0001).

**Fig. 2.**
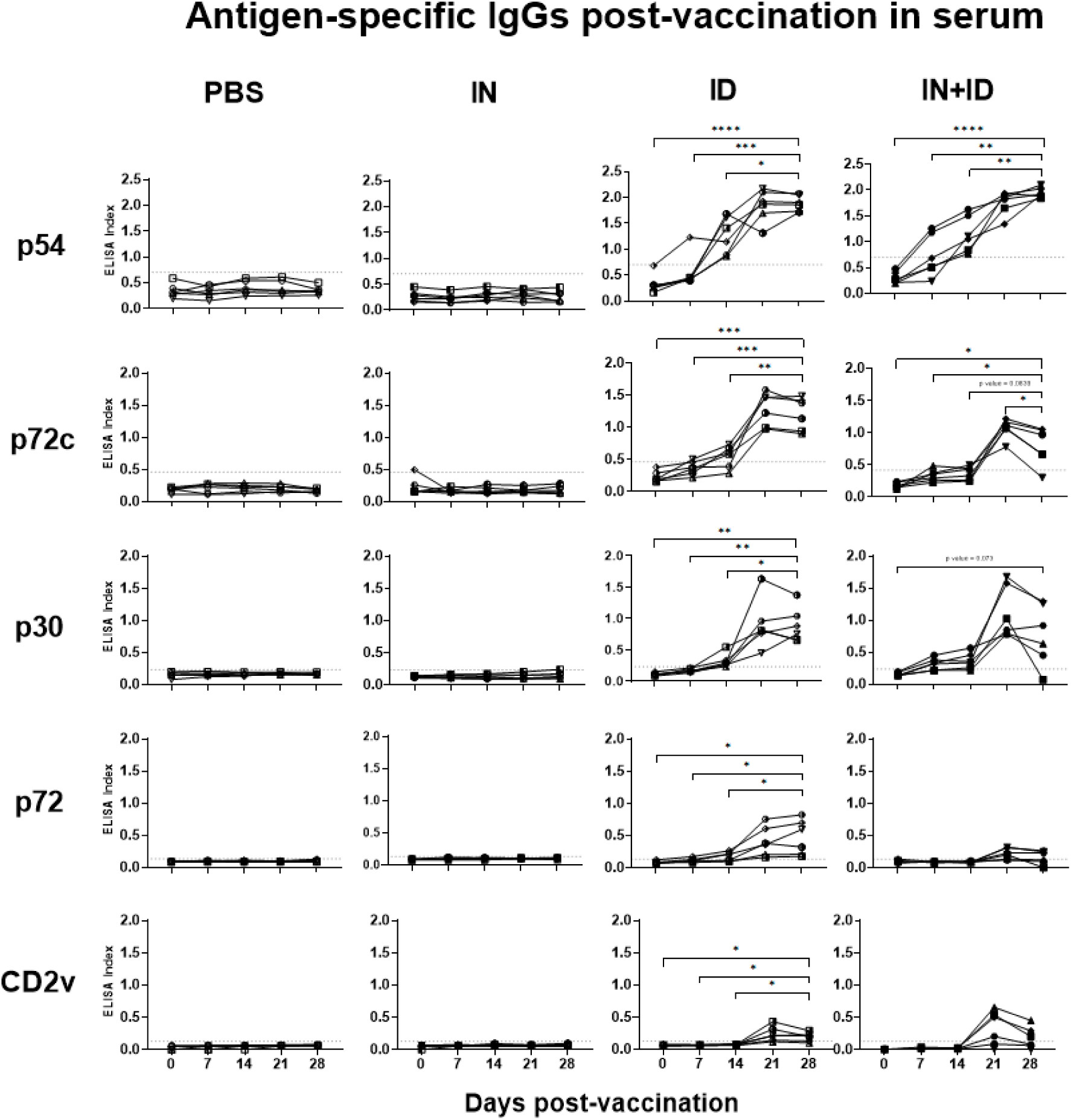
Antigen-specific IgG antibodies following vaccination with the ASFV protein cocktail. Serum IgG antibodies against p54, p72c, p30, p72, and CD2v were measured by ELISA from 0 to 28 dpv in pigs vaccinated through the ID, IN and ID+IN routes. Statistical significance was determined by one-way ANOVA (\**P* < 0.05, \*\**P* < 0.01, \*\*\**P* < 0.001 and \*\*\*\**P* < 0.0001).

Having characterised the humoral immune response, we next evaluated vaccine- induced cellular immunity. IFN-ɣ-producing cells were measured by ELISpot following *in vitro* stimulation with individual ASFV proteins at 28 dpv (Fig. 3) or with ASFV Georgia 2007/1 at 14 and 28 dpv (Fig. S4). Among the antigens tested, only p54 induced significantly higher antigen-specific IFN-ɣ-producing cells in the ID group compared with controls. Although statistical significance was not achieved, p30 and p72c showed a variable number of IFN-ɣ-producing cells from 50 to 170 SFU/10^6^ PBMCs. Interestingly, the IN group showed a trend towards significance following p54 stimulation (*P* value = 0.07), despite the absence of a detectable systemic antibody response. In contrast, stimulation with ASFV Georgia 2007/1 did not induce significant IFN-ɣ-producing T-cells in any group at either time point. The frequency of perforin- positive T-cell subsets was also evaluated by flow cytometry following stimulation with ASFV Georgia2007/1, or the proteins used for vaccination combined and individually (Fig. S5 and S6). No significant differences were detected between vaccinated and control animals at either 14 or 28 dpv. Together, these results indicate that vaccination induced only limited ASFV-specific cellular immune responses.

**Fig. 3.**
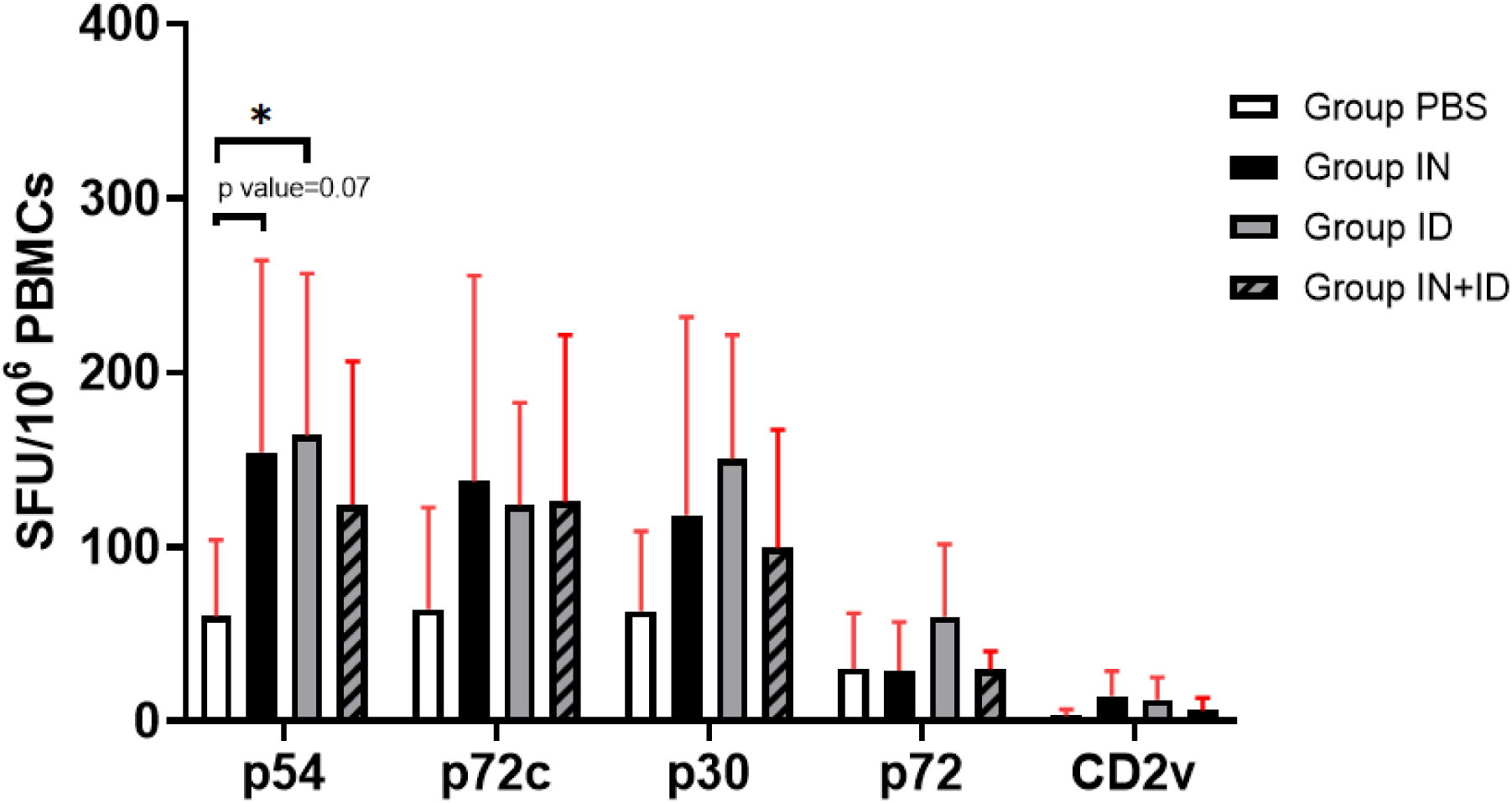
Antigen-specific IFN-ɣ-producing cells induced by vaccination. PBMCs collected at 28 dpv were stimulated *in vitro* with the individual ASFV proteins included in the vaccine cocktail (p54, p72c, p30, p72 and CD2v), and IFN-ɣ-producing cells were quantified by ELISpot. Statistical significance was determined by a two-way ANOVA (\**P* < 0.05, \*\**P* < 0.01 and \*\*\**P* < 0.001).

### 3.2. Lack of protection against virulent ASFV challenge regardless of the vaccine route

To evaluate the protective efficacy of the vaccine formulation and the influence of administration routes, pigs were challenged four weeks after the first vaccine dose with 10^5^ HAU of the highly virulent Georgia2007/1 strain administered intranasally (Fig. 1A). All animals, regardless of the experimental group, developed fever and clinical signs compatible with ASF between 4-6 days post-challenge (dpc) and reached humane endpoints at 7-8 dpc, requiring euthanasia (Fig. 4A and B). Macroscopic lesions were also comparable among groups and included enlarged haemorrhagic lymph nodes, petechiae in kidney capsule and splenomegaly, all characteristic findings of acute ASF (Fig. 4C). Although vaccinated animals appeared to display slightly less severe renal lesions than control animals, no consistent differences in clinical outcome were observed. Likewise, virus loads in blood and nasal swabs at 7 dpc were similar in all groups, reaching approximately 10^9^ and 10^7^ GEC/mL, respectively (Fig. 5). Together, these results indicate that the vaccine failed to confer protection against virulent ASFV challenge regardless of the administration route.

**Fig. 4.**
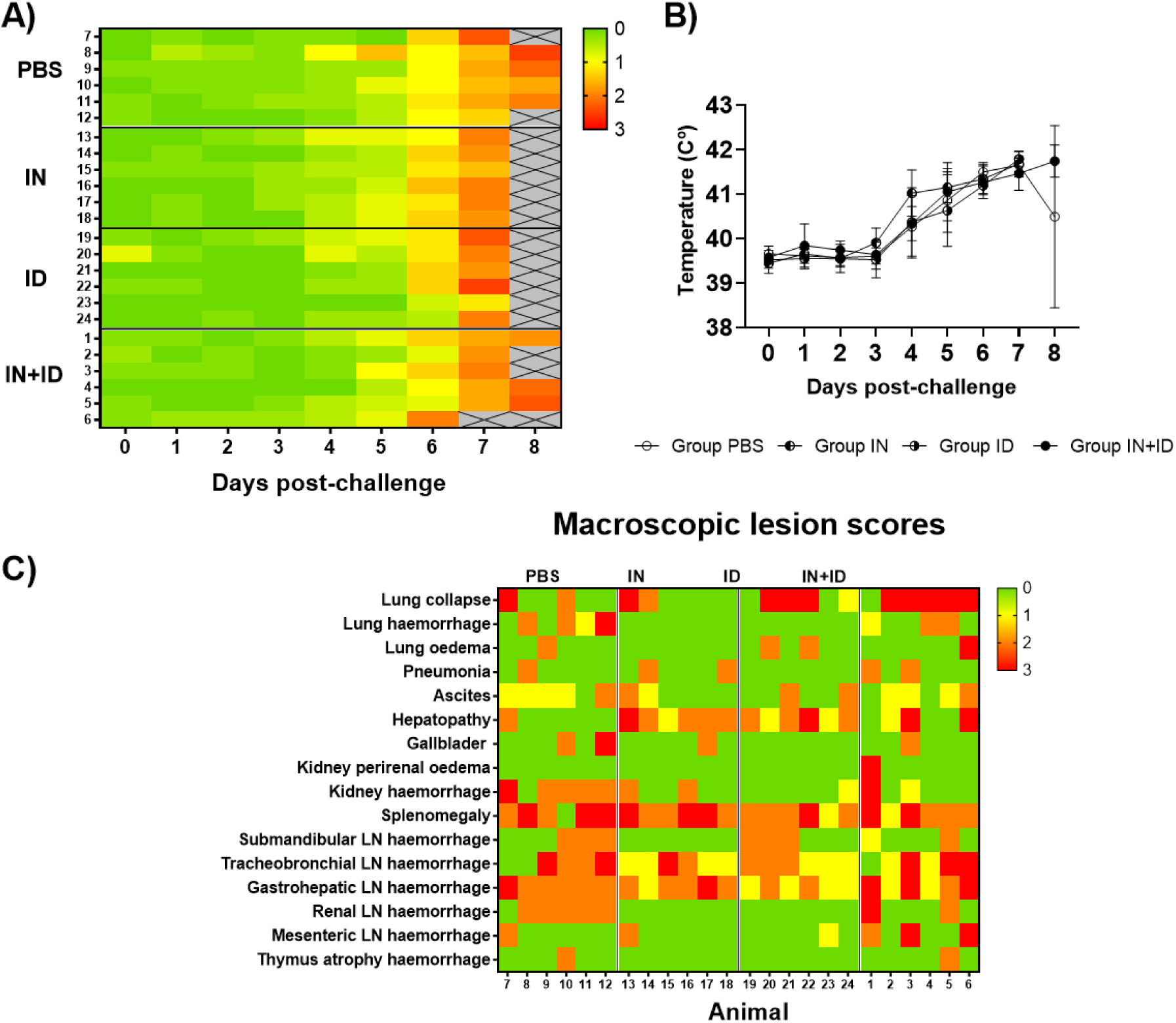
Clinical outcome and pathological findings following virulent ASFV challenge. Daily clinical scores (A) and rectal temperatures (B) after challenge. (C) Severity of macroscopic lesions associated with ASF observed at necropsy. Colours indicate lesion severity: green (absent), yellow (mild), orange (moderate) and red (severe).

**Fig. 5.**
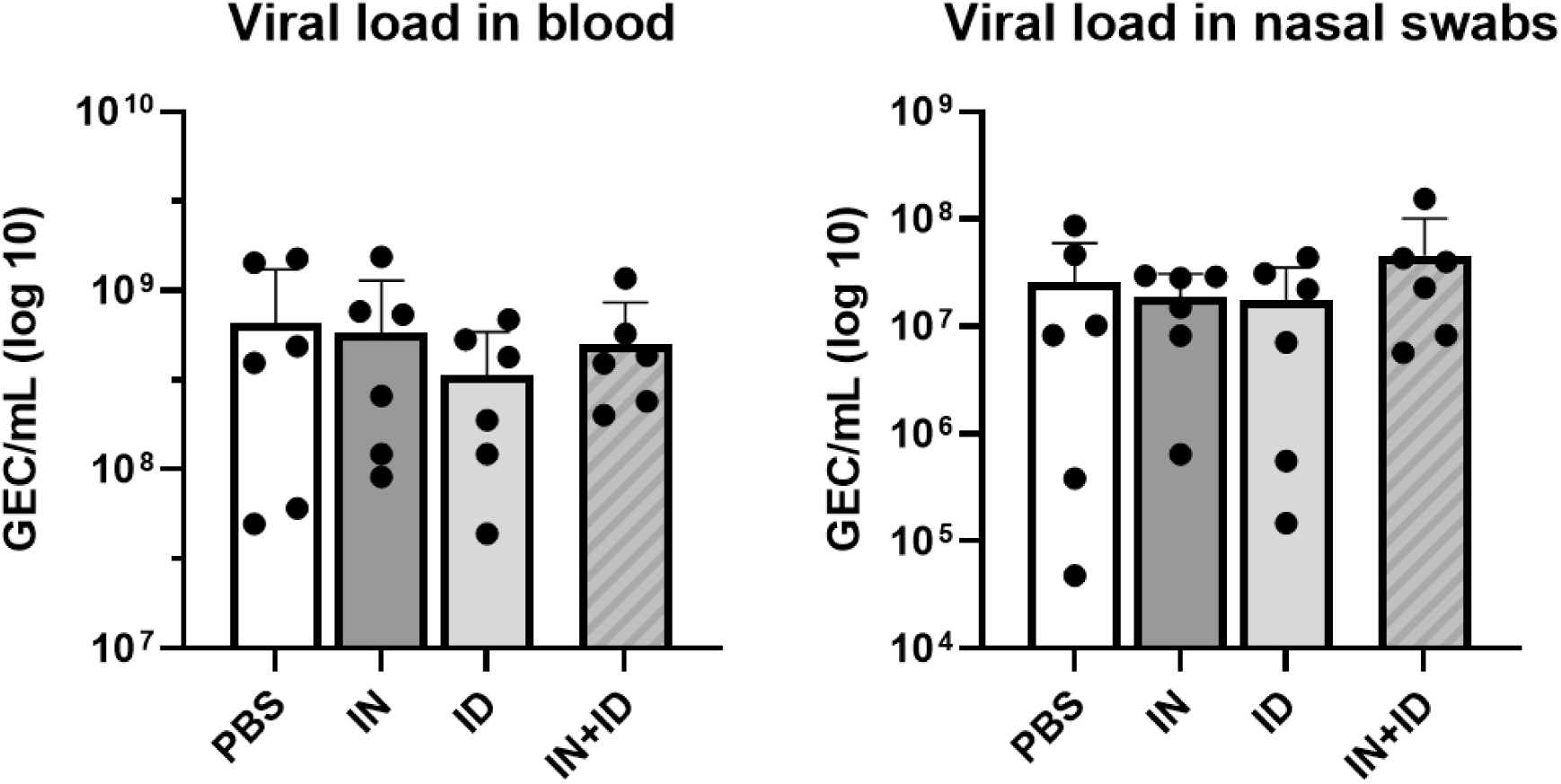
Viral loads following ASFV challenge. ASFV genome equivalent copies (GEC/ml) quantified by qPCR in blood and nasal swabs collected at 7 dpc.

### 3.3. Boost of systemic ASFV-specific humoral response following challenge in intranasally vaccinated pigs

To investigate the impact of vaccination route on recall immune responses following ASFV infection, ASFV-specific antibody responses were analysed in serum and in nasal swabs at 0 and 7 dpc. Interestingly, IgM levels increased significantly only in animals that had received IN vaccination, either alone or in combination with the ID route, whereas changes in the control and ID groups did not reach statistical significance (Fig. 6A). Likewise, ASFV-specific IgG levels increased significantly only in the IN group following challenge (Fig. 6B). Antigen-specific ELISA revealed that this increase was predominantly directed against the p54 protein (Fig. S7). In contrast, ASFV-specific and antigen-specific IgG levels decreased in animals vaccinated through the ID or ID+IN routes (Fig. 6B and S7). To evaluate the potential induction of mucosal humoral responses, we next analysed antigen-specific IgA at different times after vaccination and challenge. IgA responses against the vaccine antigens in nasal swabs were low and variable, showing that the most consistent IgA response was observed against p54 in the ID route (Fig. 7). Similarly to IgM responses, although all groups were negative for IgA antibodies in serum following vaccination, after challenge animals vaccinated through the combined ID+IN routes showed a robust boost of p54-specific IgA antibodies (Fig. 8). Together, these results suggest that IN vaccination induced immune priming that was not detected systemically prior to challenge but became evident following ASFV infection.

**Fig. 6.**
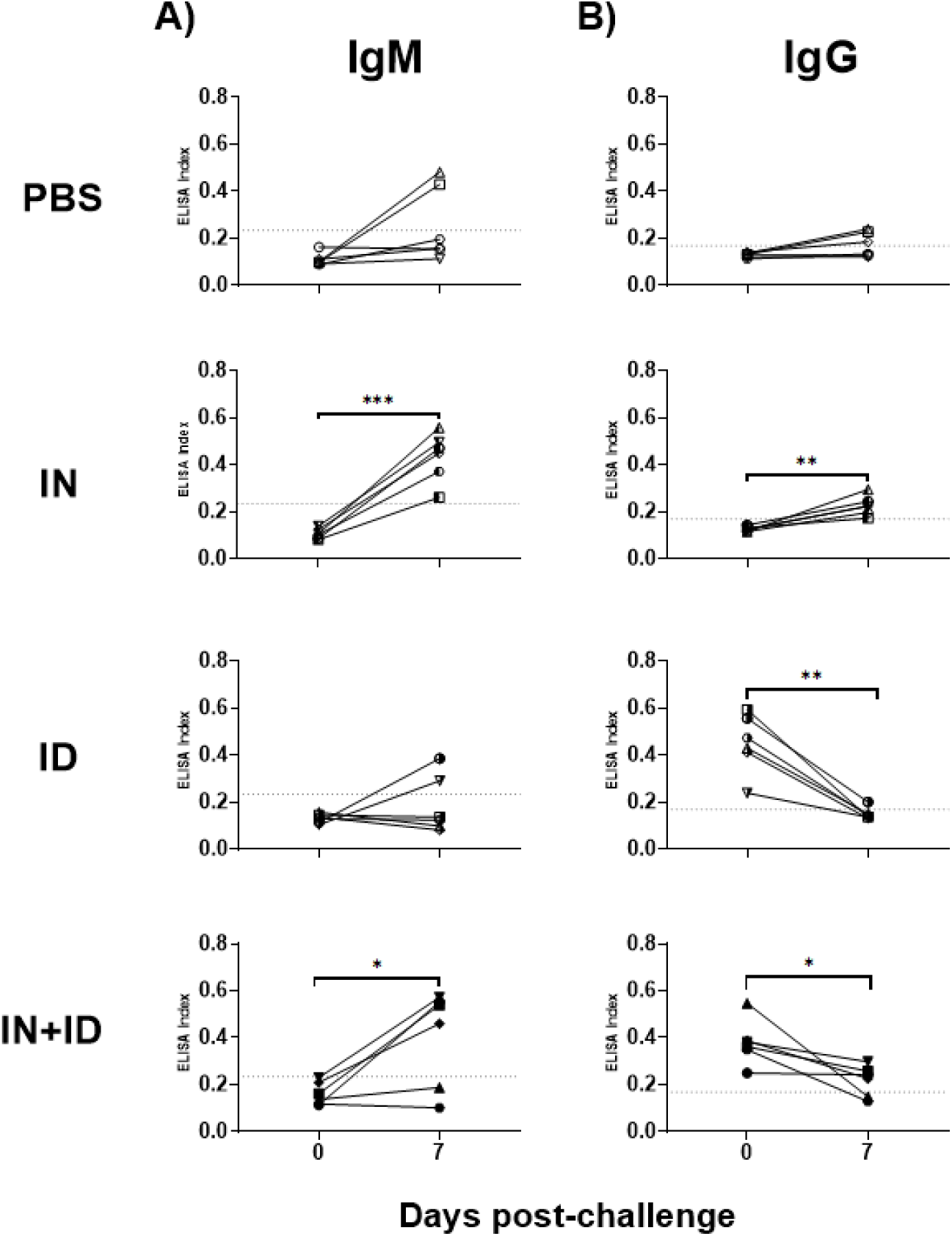
Virus-specific antibody responses following virulent ASFV challenge. ASFV-specific IgM (A) and IgG (B) antibodies in serum at 0 and 7 dpc measured by ELISA. Statistical significance was determined by a paired t-test (\**P* < 0.05, \*\**P* < 0.005 and \*\*\**P* < 0.0005).

**Fig. 7.**
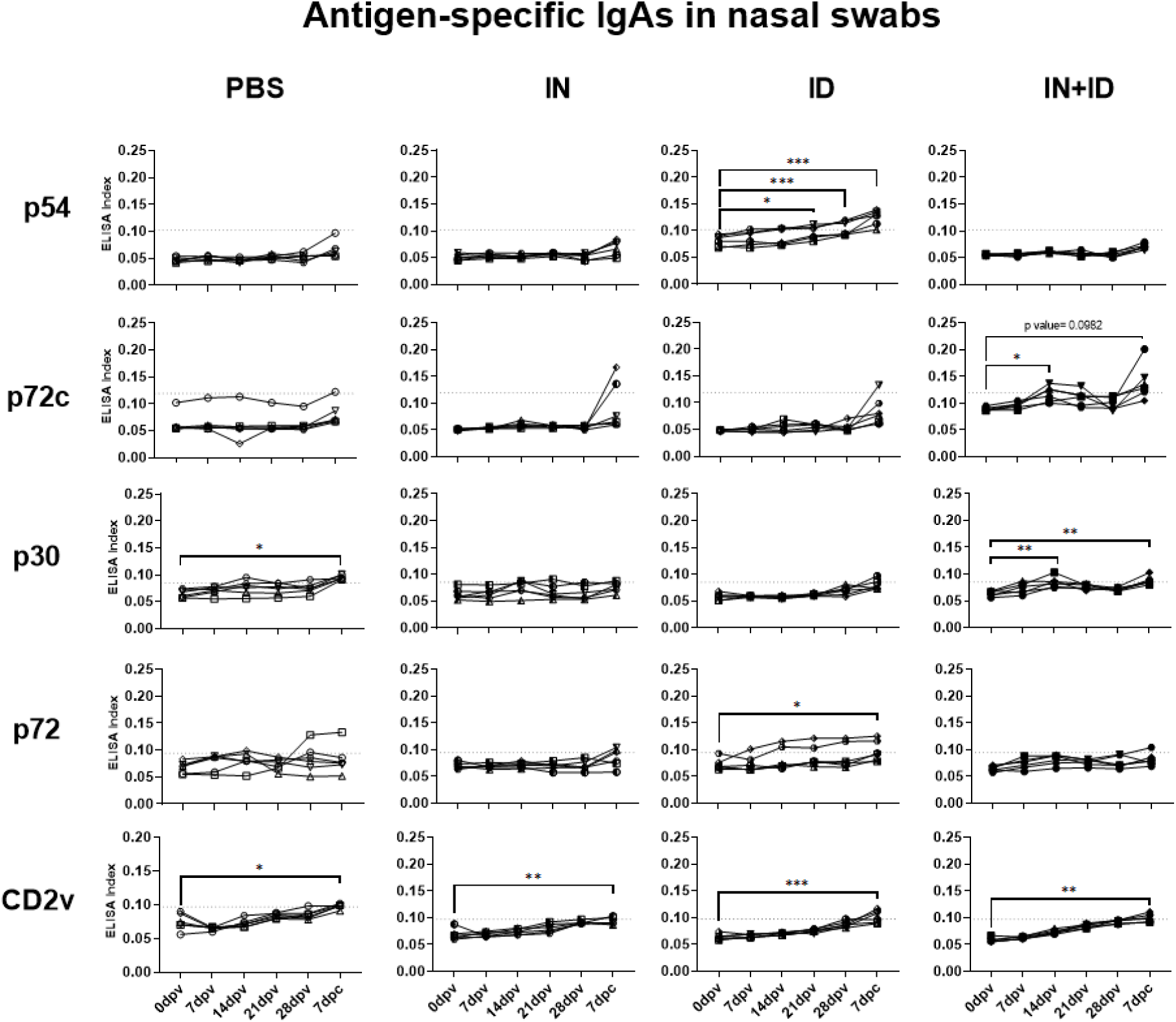
Antigen-specific IgA responses from nasal swabs throughout the experiment. IgA antibodies against p54, p72c, p30, p72, and CD2v were measured by ELISA in pigs vaccinated through the ID, IN and ID+IN routes. Statistical significance was determined by one-way ANOVA (\**P* < 0.05, \*\**P* < 0.01, \*\*\**P* < 0.001 and \*\*\*\**P* < 0.0001).

**Fig. 8.**
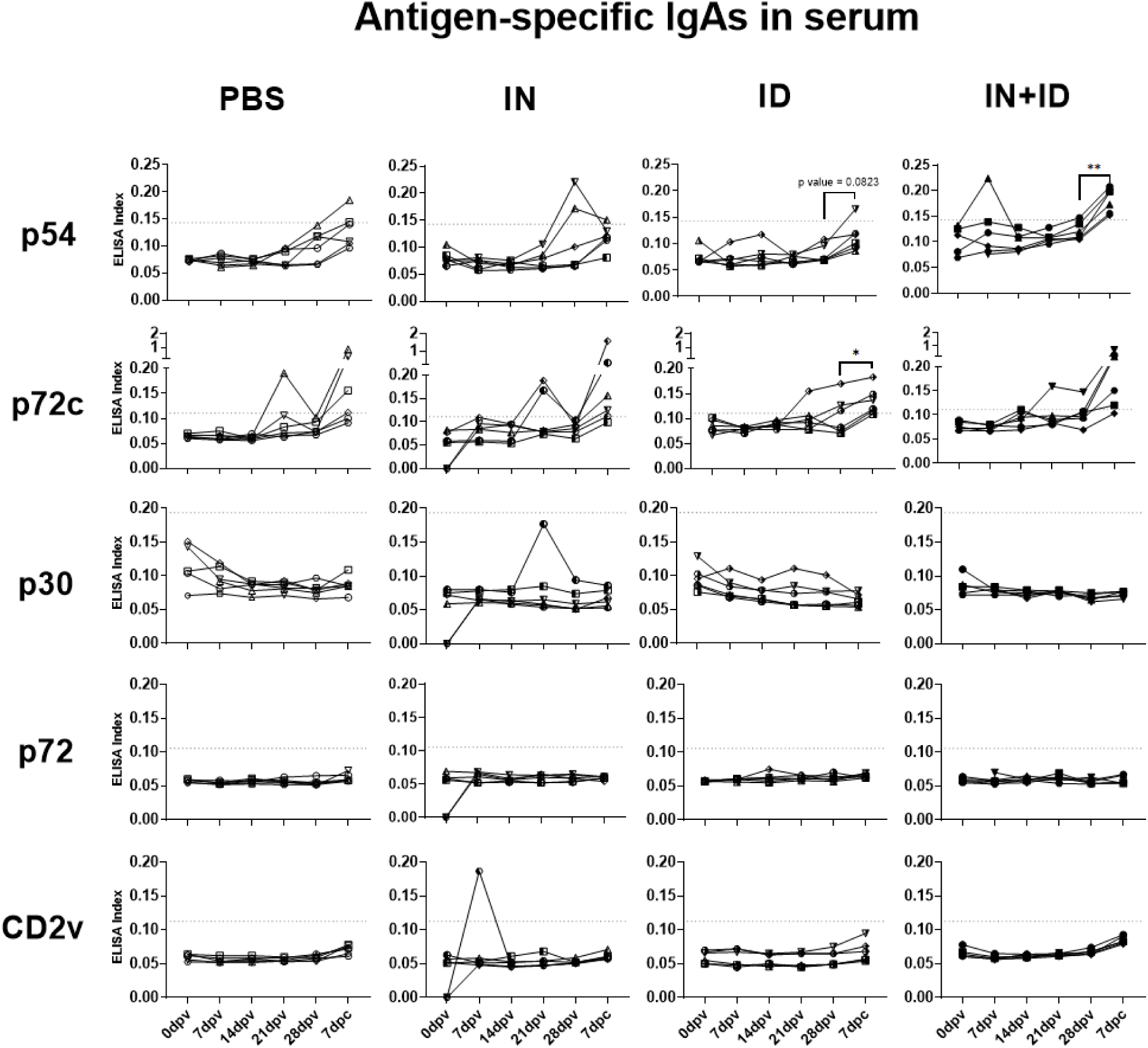
Antigen-specific serum IgA responses throughout the experiment. Serum IgA antibodies against p54, p72c, p30, p72, and CD2v were measured by ELISA in pigs vaccinated through the ID, IN and ID+IN routes. Statistical significance was determined by one-way ANOVA (\**P* < 0.05, \*\**P* < 0.01, \*\*\**P* < 0.001 and \*\*\*\**P* < 0.0001).

## 4. Discussion

Most ASFV subunit vaccine studies have focused on identifying protective antigens and immune correlates of protection [41]. However, vaccine efficacy is also critically influenced by antigen delivery systems as well as administration routes. In this context, the results of the present study provide further evidence that the same antigen combination can generate markedly different immune outcomes depending on how it is delivered. The five ASFV proteins used in our vaccine formulation were selected from the study performed by Liu et al., where the same combination fully protected animals against ASF when delivered in an adenovirus-vectored platform [4]. In contrast, the secretory granule-based formulation evaluated here failed to induce broad immune responses and did not confer protection following virulent ASFV challenge. Together, these findings indicate that antigen selection alone is insufficient to predict vaccine efficacy and further highlight the importance of delivery systems in shaping protective immunity.

Viral vector-based vaccines, such as adenovirus- and recombinant PRV-based platforms, have shown promising results in experimental swine vaccination studies [42]. Their ability to promote intracellular antigen expression facilitates presentation through both SLA-I and SLA-II pathways, thereby efficiently inducing cellular immune responses [6], [41], [42], [43]. In contrast, despite the inherent immunostimulatory properties previously described for the secretory granule platform [33], the microparticulated vaccine evaluated here induced only limited ASFV-specific cellular immunity. This may represent an important limitation for protection against ASFV, as cellular immune responses, and particularly cytotoxic T cells, are considered critical components of protective immunity [13], [44]. Indeed, our results further reinforce the growing evidence that robust humoral responses alone are insufficient to confer protection against ASFV [45]. Nevertheless, despite the promising performance of viral vectors, recombinant protein-based vaccines remain an attractive alternative due to their favourable safety profile, manufacturing flexibility and compatibility with repeated administration [46]. In this context, the secretory granule platform will likely require further optimization to enhance the induction of cellular immune responses, potentially through the incorporation of adjuvants [6], [42]. Finally, direct comparisons between the present study and that of Liu et al. should be interpreted with caution, as protection reported in the latter was associated with a natural field exposure rather than a standardized experimental challenge.

Whether the combination of p30, p54, p72, p72c and CD2v is sufficient to confer robust protection against ASFV remains unresolved. Before the study from Liu et al. [4], several of these antigens were included in different subunit, DNA and viral-vectored vaccine formulations resulting in inconsistent efficacy results in terms of immunogenicity and protection conferred, ranging from lack of protection to only partial protection [2], [41], [45], [47], [48], [49]. A notable example is the work by Jancovich et al., in which a pool of 35 ASFV antigens, including p30, p54, p72, p72c and CD2v, using a DNA prime and recombinant vaccinia virus boost approach, induced both humoral and cellular immune responses but failed to confer clinical protection [50]. In contrast, Goatley et al. showed that the removal of p54 and p72c from a protective viral-vectored cocktail including eight antigens abolished protection, suggesting an important contribution of these two antigens to vaccine-induced protective immunity [3]. Consistently, in our study p54 consistently emerged as one of the most immunogenic components of the vaccine formulation, inducing both antibody and cellular responses and representing the main target of the accelerated post-challenge antibody boost observed in intranasally vaccinated animals. More recently, Portugal et al. reported that antigen combinations capable of inducing protection against a genotype I challenge did not protect when tailored against a genotype II virus, further highlighting the complexity of identifying pan-protective ASFV antigens [1]. Overall, these observations suggest that the vaccine platform and vaccination strategy used to test ASFV antigens are critical factors influencing their protective potential. Thus, the complete protection reported by Liu et al. may result not only from the antigen combination selected, but also from the capacity of the adenoviral platform and the combined intramuscular and intranasal administrations used to induce protective immunity.

In contrast to antigen selection and vaccine delivery systems, the importance of the administration route has often been neglected. In this regard, the complete protection reported by Liu et al. may have been influenced by the combined use of the intramuscular and intranasal vaccine delivery [4]. However, as the specific contribution of each route was not evaluated, the relative role of mucosal and systemic immunisation in protection could not be determined. Here, we evaluated whether intranasal administration of the same ASFV antigens could influence vaccine-induced immune responses and contribute to protection against ASFV. Although intranasal vaccination induced evidence of immune priming, as reflected by the accelerated IgA, IgG and IgM responses observed after challenge, the secretory granule-based formulation evaluated in this study failed to induce a robust cellular immunity and did not confer protection. Therefore, the capacity of mucosal immunisation to contribute to ASFV protection remains unresolved. Further studies using alternative subunit or live attenuated vaccine strategies are needed to clarify the role of mucosal immunity in ASFV vaccine efficacy. Importantly, such studies should avoid relying exclusively on intramuscular challenge models, as bypassing the natural routes of viral entry may limit the ability to elucidate the contributions of early mucosal immune responses to protection against a natural infection.

## References

[1] R. Portugal et al., ‘Six adenoviral vectored African swine fever virus genes protect against fatal disease caused by genotype I challenge’, J. Virol., pp. e00622–24, Jul. 2024, doi: 10.1128/jvi.00622-24.

[2] L. Dong et al., ‘Recombinant vaccinia vectored ASFV vaccine enhances swine survival against genotype II challenge’, Npj Vaccines, vol. 11, no. 1, p. 58, Jan. 2026, doi: 10.1038/s41541-026-01377-0.

[3] L. C. Goatley et al., ‘A Pool of Eight Virally Vectored African Swine Fever Antigens Protect Pigs against Fatal Disease’, Vaccines, vol. 8, no. 2, p. 234, May 2020, doi: 10.3390/vaccines8020234.

[4] W. Liu et al., ‘A new vaccination regimen using adenovirus-vectored vaccine confers effective protection against African swine fever virus in swine’, Emerg. Microbes Infect., vol. 12, no. 2, p. 2233643, Dec. 2023, doi: 10.1080/22221751.2023.2233643.

[5] R. Kumar et al., ‘Live-vectored antigen cocktail confers protection against African swine fever virus (ASFV) Georgia 2007/1 challenge’, Npj Vaccines, Feb. 2026, doi: 10.1038/s41541-026-01399-8.

[6] Q. Wu et al., ‘Advancement in the Antigenic Epitopes and Vaccine Adjuvants of African Swine Fever Virus’, Pathogens, vol. 13, no. 8, p. 706, Aug. 2024, doi: 10.3390/pathogens13080706.

[7] L. Zhang, W. Wang, and S. Wang, ‘Effect of vaccine administration modality on immunogenicity and efficacy’, Expert Rev. Vaccines, vol. 14, no. 11, pp. 1509–1523, Nov. 2015, doi: 10.1586/14760584.2015.1081067.

[8] M. Tozuka et al., ‘Efficient antigen delivery to the draining lymph nodes is a key component in the immunogenic pathway of the intradermal vaccine’, J. Dermatol. Sci., vol. 82, no. 1, pp. 38–45, Apr. 2016, doi: 10.1016/j.jdermsci.2015.11.008.

[9] N. Romani, V. Flacher, C. H. Tripp, F. Sparber, S. Ebner, and P. Stoitzner, ‘Targeting Skin Dendritic Cells to Improve Intradermal Vaccination’, in Intradermal Immunization, M. B. M. Teunissen, Ed., Berlin, Heidelberg: Springer, 2012, pp. 113–138. doi: 10.1007/82_2010_118.

[10] S. Fan et al., ‘Role of innate lymphoid cells and dendritic cells in intradermal immunization of the enterovirus antigen’, Npj Vaccines, vol. 4, no. 1, p. 14, Mar. 2019, doi: 10.1038/s41541-019-0108-6.

[11] A. C.-Y. Lee et al., ‘Intradermal vaccination of live attenuated influenza vaccine protects mice against homologous and heterologous influenza challenges’, Npj Vaccines, vol. 6, no. 1, p. 95, Aug. 2021, doi: 10.1038/s41541-021-00359-8.

[12] J.-B. Le Luduec et al., ‘Intradermal vaccination with un-adjuvanted sub- unit vaccines triggers skin innate immunity and confers protective respiratory immunity in domestic swine’, Vaccine, vol. 34, no. 7, pp. 914–922, Feb. 2016, doi: 10.1016/j.vaccine.2015.12.058.

[13] M. Pedrera et al., ‘Characterization of the Protective Cellular Immune Response in Pigs Immunized Intradermally with the Live Attenuated African Swine Fever Virus (ASFV) Lv17/WB/Rie1’, Vaccines, vol. 12, no. 4, p. 443, Apr. 2024, doi: 10.3390/vaccines12040443.

[14] S. Lee et al., ‘Intradermal Inoculation of Inactivated Foot-and-Mouth Disease Vaccine Induced Effective Immune Responses Comparable to Conventional Intramuscular Injection in Pigs’, Vaccines, vol. 12, no. 2, p. 190, Feb. 2024, doi: 10.3390/vaccines12020190.

[15] M. Salman et al., ‘Intradermal needle-free injection prevents African Swine Fever transmission, while intramuscular needle injection does not’, Sci. Rep., vol. 13, no. 1, p. 4600, Mar. 2023, doi: 10.1038/s41598-023-31199-2.

[16] L. A. Morici and J. B. McLachlan, ‘Non-mucosal vaccination strategies to enhance mucosal immunity’, Vaccine Insights, vol. 2, no. 6, pp. 229–236, Jul. 2023, doi: 10.18609/vac.2023.034.

[17] J. R. Raha et al., ‘Intranasal vaccination with multi-neuraminidase and M2e virus-like particle vaccine results in greater mucosal immunity and protection against influenza than intramuscular injection’, Vaccine, vol. 57, p. 127206, May 2025, doi: 10.1016/j.vaccine.2025.127206.

[18] C. Singh et al., ‘Phase III Pivotal comparative clinical trial of intranasal (iNCOVACC) and intramuscular COVID 19 vaccine (Covaxin®)’, Npj Vaccines, vol. 8, no. 1, p. 125, Aug. 2023, doi: 10.1038/s41541-023-00717-8.

[19] D. R. Braucher et al., ‘Intranasal Vaccination with Replication-Defective Adenovirus Type 5 Encoding Influenza Virus Hemagglutinin Elicits Protective Immunity to Homologous Challenge and Partial Protection to Heterologous Challenge in Pigs’, Clin. Vaccine Immunol., vol. 19, no. 11, pp. 1722–1729, Nov. 2012, doi: 10.1128/CVI.00315-12.

[20] R. Avanthay et al., ‘Evaluation of a novel intramuscular prime/intranasal boost vaccination strategy against influenza in the pig model’, PLOS Pathog., vol. 20, no. 8, p. e1012393, Aug. 2024, doi: 10.1371/journal.ppat.1012393.

[21] S. Dhakal et al., ‘Mucosal Immunity and Protective Efficacy of Intranasal Inactivated Influenza Vaccine Is Improved by Chitosan Nanoparticle Delivery in Pigs’, Front. Immunol., vol. 9, May 2018, doi: 10.3389/fimmu.2018.00934.

[22] P. Chaikhumwang, A. Madapong, K. Saeng-chuto, D. Nilubol, and A. Tantituvanont, ‘Intranasal delivery of inactivated PRRSV loaded cationic nanoparticles coupled with enterotoxin subunit B induces PRRSV-specific immune responses in pigs’, Sci. Rep., vol. 12, no. 1, p. 3725, Mar. 2022, doi: 10.1038/s41598-022-07680-9.

[23] E. Mateu et al., ‘Efficacy of an intranasally administered live attenuated PRRSV-2 vaccine against challenge with a highly virulent PRRSV-1 strain’, Front. Vet. Sci., vol. 12, p. 1619052, Aug. 2025, doi: 10.3389/fvets.2025.1619052.

[24] J. F. Hernandez-Franco et al., ‘Intradermal Vaccination against Influenza with a STING-Targeted Nanoparticle Combination Adjuvant Induces Superior Cross-Protective Humoral Immunity in Swine Compared with Intranasal and Intramuscular Immunization’, Vaccines, vol. 11, no. 11, p. 1699, Nov. 2023, doi: 10.3390/vaccines11111699.

[25] A. Prapattanakullawong et al., ‘Comparative evaluation of PRRSV MLV vaccine efficacy administered via intranasal and intramuscular routes in early age of piglets’, Vaccine, vol. 65, p. 127822, Oct. 2025, doi: 10.1016/j.vaccine.2025.127822.

[26] Z. Zhang et al., ‘Comparative immunogenicity analysis of intradermal versus intramuscular immunization with a recombinant human adenovirus type 5 vaccine against Ebola virus’, Front. Immunol., vol. 13, p. 963049, Aug. 2022, doi: 10.3389/fimmu.2022.963049.

[27] P. J. Sánchez-Cordón et al., ‘Evaluation of protection induced by immunisation of domestic pigs with deletion mutant African swine fever virus BeninΔMGF by different doses and routes’, Vaccine, vol. 36, no. 5, pp. 707–715, Jan. 2018, doi: 10.1016/j.vaccine.2017.12.030.

[28] O. Bourry et al., ‘Oronasal or Intramuscular Immunization with a Thermo- Attenuated ASFV Strain Provides Full Clinical Protection against Georgia 2007/1 Challenge’, Viruses, vol. 14, no. 12, p. 2777, Dec. 2022, doi: 10.3390/v14122777.

[29] M. V. Borca et al., ‘ASFV-G-ΔI177L as an Effective Oral Nasal Vaccine against the Eurasia Strain of Africa Swine Fever’, Viruses, vol. 13, no. 5, p. 765, May 2021, doi: 10.3390/v13050765.

[30] M. Vlasov et al., ‘Administration Routes and Doses of the Attenuated African Swine Fever Virus Strain PSA-1NH Influence Cross-Protection of Pigs against Heterologous Challenge’, Animals, vol. 14, no. 9, p. 1277, Jan. 2024, doi: 10.3390/ani14091277.

[31] P. J. Sánchez-Cordón et al., ‘Different routes and doses influence protection in pigs immunised with the naturally attenuated African swine fever virus isolate OURT88/3’, Antiviral Res., vol. 138, pp. 1–8, Feb. 2017, doi: 10.1016/j.antiviral.2016.11.021.

[32] E. Parladé et al., ‘Subcutaneous administration of an endocrine-mimetic, slow-release protein material reduces the severity of SARS-CoV-2 infection’, J. Drug Deliv. Sci. Technol., vol. 107, p. 106813, May 2025, doi: 10.1016/j.jddst.2025.106813.

[33] L. Bosch-Camós et al., ‘Nanoparticle-Based Secretory Granules Induce a Specific and Long-Lasting Immune Response through Prolonged Antigen Release’, Nanomaterials, vol. 14, no. 5, p. 435, Feb. 2024, doi: 10.3390/nano14050435.

[34] C. L. Netherton, Ed., African Swine Fever Virus: Methods and Protocols, vol. 2503. in Methods in Molecular Biology, vol. 2503. New York, NY: Springer US, 2022. doi: 10.1007/978-1-0716-2333-6.

[35] J. M. Argilaguet et al., ‘DNA Vaccination Partially Protects against African Swine Fever Virus Lethal Challenge in the Absence of Antibodies’, PLOS ONE, vol. 7, no. 9, p. e40942, Sep. 2012, doi: 10.1371/journal.pone.0040942.

[36] L. J. Reed and H. Muench, ‘A SIMPLE METHOD OF ESTIMATING FIFTY PER CENT ENDPOINTS’, Am. J. Epidemiol., vol. 27, no. 3, pp. 493–497, May 1938, doi: 10.1093/oxfordjournals.aje.a118408.

[37] I. Galindo-Cardiel et al., ‘Standardization of pathological investigations in the framework of experimental ASFV infections’, Virus Res., vol. 173, no. 1, pp. 180–190, Apr. 2013, doi: 10.1016/j.virusres.2012.12.018.

[38] A. Lacasta et al., ‘Expression Library Immunization Can Confer Protection against Lethal Challenge with African Swine Fever Virus’, J. Virol., vol. 88, no. 22, pp. 13322–13332, Nov. 2014, doi: 10.1128/JVI.01893-14.

[39] C. Gallardo et al., ‘Comparative evaluation of novel African swine fever virus (ASF) antibody detection techniques derived from specific ASF viral genotypes with the OIE internationally prescribed serological tests’, Vet. Microbiol., vol. 162, no. 1, pp. 32–43, Feb. 2013, doi: 10.1016/j.vetmic.2012.08.011.

[40] I. Díaz and E. Mateu, ‘Use of ELISPOT and ELISA to evaluate IFN- gamma, IL-10 and IL-4 responses in conventional pigs’, Vet. Immunol. Immunopathol., vol. 106, no. 1–2, pp. 107–112, Jun. 2005, doi: 10.1016/j.vetimm.2005.01.005.

[41] P. Y. L. Tng, L. Al-Adwani, L. Goatley, R. Portugal, A. Rathakrishnan, and C. L. Netherton, ‘Early disruption of the innate-adaptive immune axis in vivo after infection with virulent Georgia 2007/1 ASFV’, Discov. Immunol., vol. 4, no. 1, p. kyaf014, Jan. 2025, doi: 10.1093/discim/kyaf014.

[42] Y. Zhang et al., ‘ASFV subunit vaccines: Strategies and prospects for future development’, Microb. Pathog., vol. 197, p. 107063, Dec. 2024, doi: 10.1016/j.micpath.2024.107063.

[43] J. M. Argilaguet et al., ‘Enhancing DNA immunization by targeting ASFV antigens to SLA-II bearing cells’, Vaccine, vol. 29, no. 33, pp. 5379–5385, Jul. 2011, doi: 10.1016/j.vaccine.2011.05.084.

[44] D. Marín-Moraleda et al., ‘Distinct cytotoxic cell subsets underlie protective and non-protective immunity to African swine fever virus’, Emerg. Microbes Infect., vol. 15, no. 1, p. 2685920, Aug. 2026, doi: 10.1080/22221751.2026.2685920.

[45] J. G. Neilan, L. Zsak, Z. Lu, T. G. Burrage, G. F. Kutish, and D. L. Rock, ‘Neutralizing antibodies to African swine fever virus proteins p30, p54, and p72 are not sufficient for antibody-mediated protection’, Virology, vol. 319, no. 2, pp. 337–342, Feb. 2004, doi: 10.1016/j.virol.2003.11.011.

[46] J. Tang, M. A. Amin, and J. L. Campian, ‘Past, Present, and Future of Viral Vector Vaccine Platforms: A Comprehensive Review’, Vaccines, vol. 13, no. 5, p. 524, May 2025, doi: 10.3390/vaccines13050524.

[47] J. M. Argilaguet et al., ‘DNA Vaccination Partially Protects against African Swine Fever Virus Lethal Challenge in the Absence of Antibodies’, PLOS ONE, vol. 7, no. 9, p. e40942, Sep. 2012, doi: 10.1371/journal.pone.0040942.

[48] M. D. Zajac et al., ‘Immunization of pigs with replication-incompetent adenovirus-vectored African swine fever virus multi-antigens induced humoral immune responses but no protection following contact challenge’, Front. Vet. Sci., vol. 10, p. 1208275, Jun. 2023, doi: 10.3389/fvets.2023.1208275.

[49] S.-Y. Sunwoo et al., ‘DNA-Protein Vaccination Strategy Does Not Protect from Challenge with African Swine Fever Virus Armenia 2007 Strain’, Vaccines, vol. 7, no. 1, p. 12, Mar. 2019, doi: 10.3390/vaccines7010012.

[50] J. K. Jancovich et al., ‘Immunization of Pigs by DNA Prime and Recombinant Vaccinia Virus Boost To Identify and Rank African Swine Fever Virus Immunogenic and Protective Proteins’, J. Virol., vol. 92, no. 8, pp. e02219–17, Apr. 2018, doi: 10.1128/JVI.02219-17.

