## Supplementary material for "Intradermal and intranasal vaccination with microparticulated ASFV proteins induces distinct immune profiles in pigs"

1 Unitat Mixta d'Investigació IRTA-UAB en Sanitat Animal, Centre de Recerca en Sanitat Animal (CReSA), Campus de la Universitat Autònoma de Barcelona (UAB), 08193 Bellaterra, Catalonia, Spain.
2 Institut de Recerca i Tecnologia Agroalimentàries (IRTA), Programa de Sanitat Animal, Centre de Recerca en Sanitat Animal (CReSA), Campus de la Universitat Autònoma de Barcelona (UAB), 08193 Bellaterra, Spain.
3 WOAH Collaborating Centre for the Research and Control of Emerging and Re-Emerging Swine Diseases in Europe (IRTA-CReSA), 08193 Bellaterra, Spain.
4 Institut de Biotecnologia i de Biomedicina (IBB) and Departament de Genètica i de Microbiologia, Universitat Autonoma` de Barcelona, Barcelona 08193, Spain.
5 IRTA, Ruminant Production, Torre Marimon, 08140, Caldes de Montbui, Catalonia, Spain.
6 Departament de Sanitat i d'Anatomia Animals. Facultat de Veterinària. Universitat Autònoma de Barcelona (UAB). Bellaterra.
* *

∮these authors equally contributed to this work.
*co-corresponding authors


### SUPPLEMENTARY MATERIAL

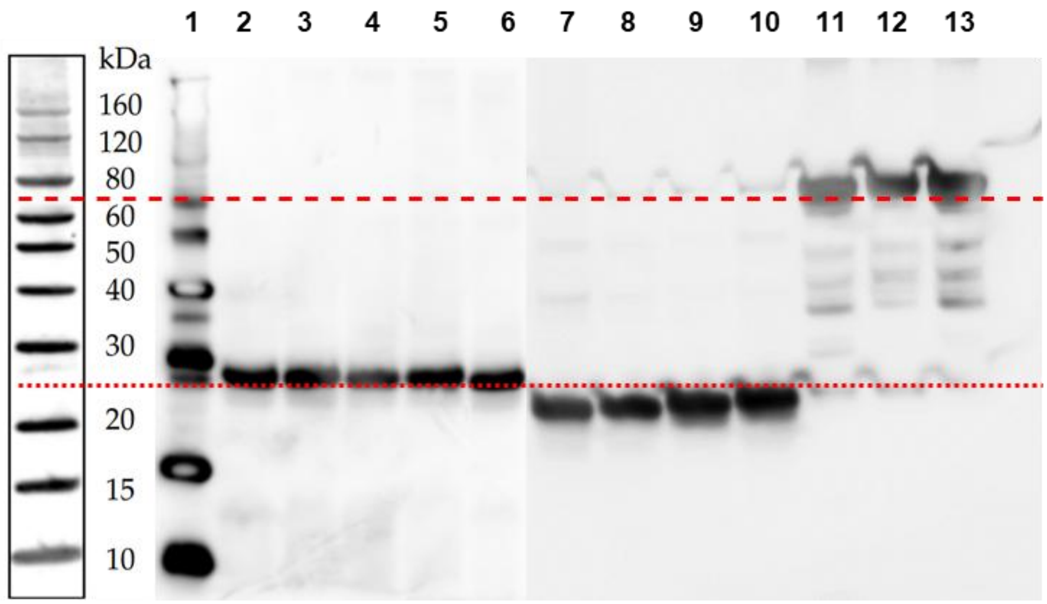


**Fig. S1.** WB of in-house produced proteins: p30 (column 2-6, around 25 kDa), p54 (column 7-10, around 25 kDa) and p72c (column 11-13, around 80 kDa). BenchMark™ His-tagged Protein Standard is in column 1, and the reference ladder is on the left of the figure.

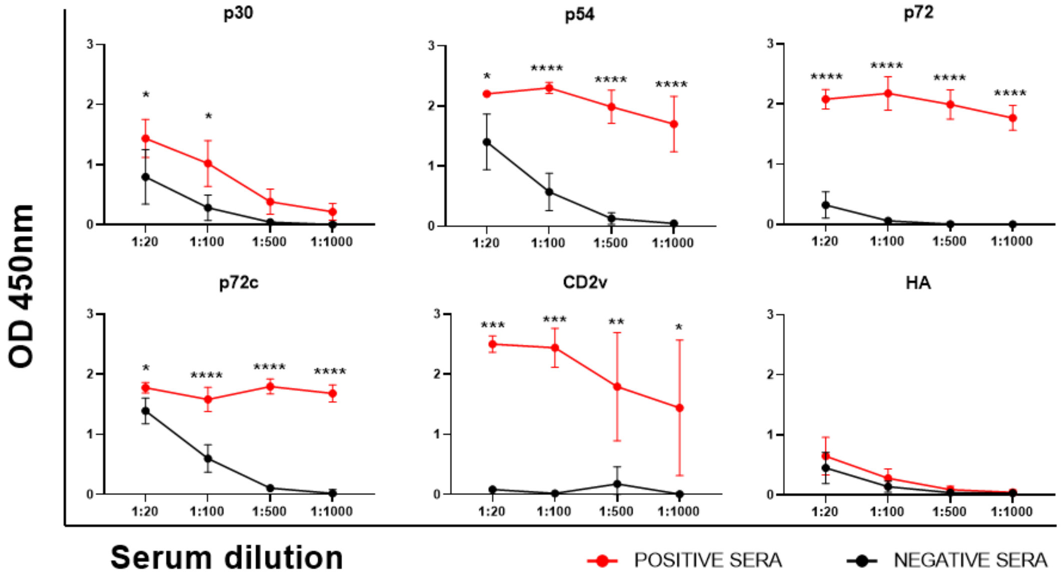


**Fig. S2.** Antigenicity of ASFV proteins included in the vaccine formulation. Recombinant p30, p54, p72, p72c and CD2v proteins were recognized by IgG antibodies present in immune sera from pigs vaccinated with Ba71ΔCD2v and subsequently challenged with Georgia2007/1. Sera from naïve pigs and recombinant H5N1 hemagglutinin (HA) were used as negative controls for antibody binding and antigen specificity, respectively. Statistical significance was determined by a two-way ANOVA (*$P < 0.05$, **$P < 0.01$, ***$P < 0.001$ and ****$P < 0.0001$).

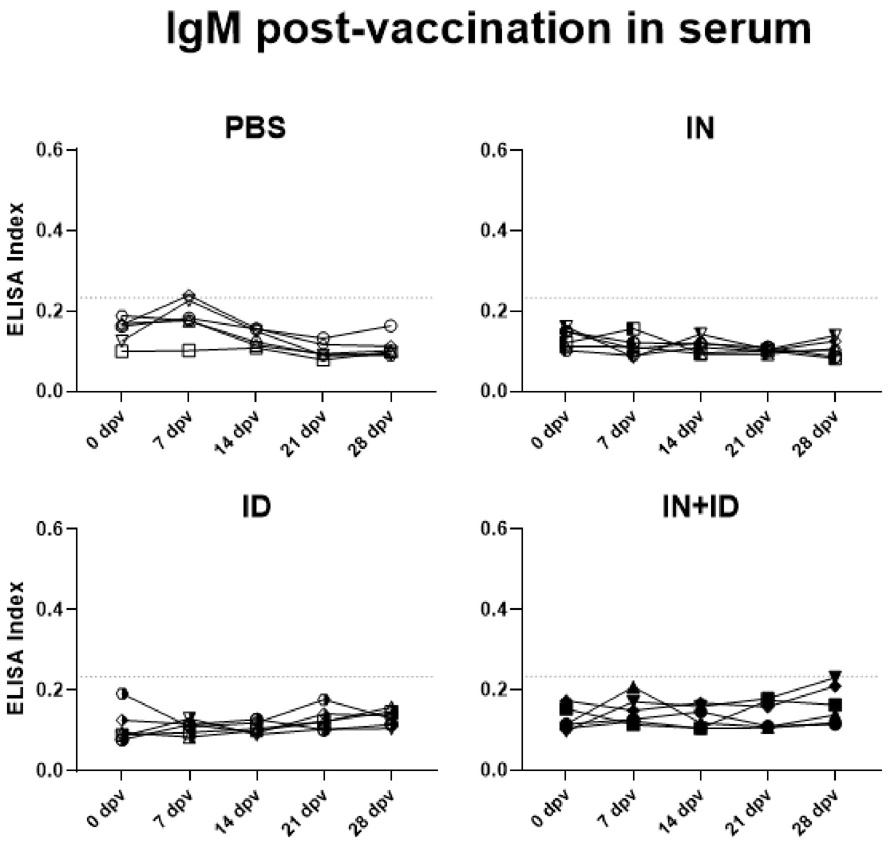


**Fig. S3.** ASFV-specific IgM antibodies following vaccination. Levels of serum virus-specific IgM antibodies were measured by ELISA from 0 to 28 dpv in pigs vaccinated via the ID, IN or ID+IN routes. PBS-inoculated pigs were used as controls. Statistical significance was determined by one-way ANOVA (**P* < 0.05, ***P* < 0.01 and ****P* < 0.001).

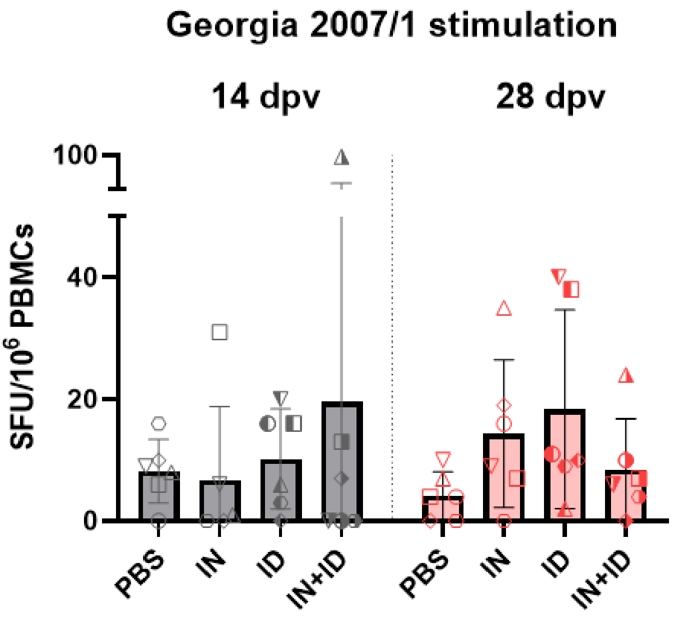


**Fig. S4.** ASFV-specific IFN-ɣ-producing T-cells following *in vitro* stimulation with ASFV Georgia 2007/1. PBMCs collected at 14 and 28 dpv (0 dpc) were stimulated with ASFV Georgia 2007/1 and IFN-ɣ-producing cells were quantified by ELISpot. Results are shown as spot-forming units (SFU) per $10^6$ PBMCs.

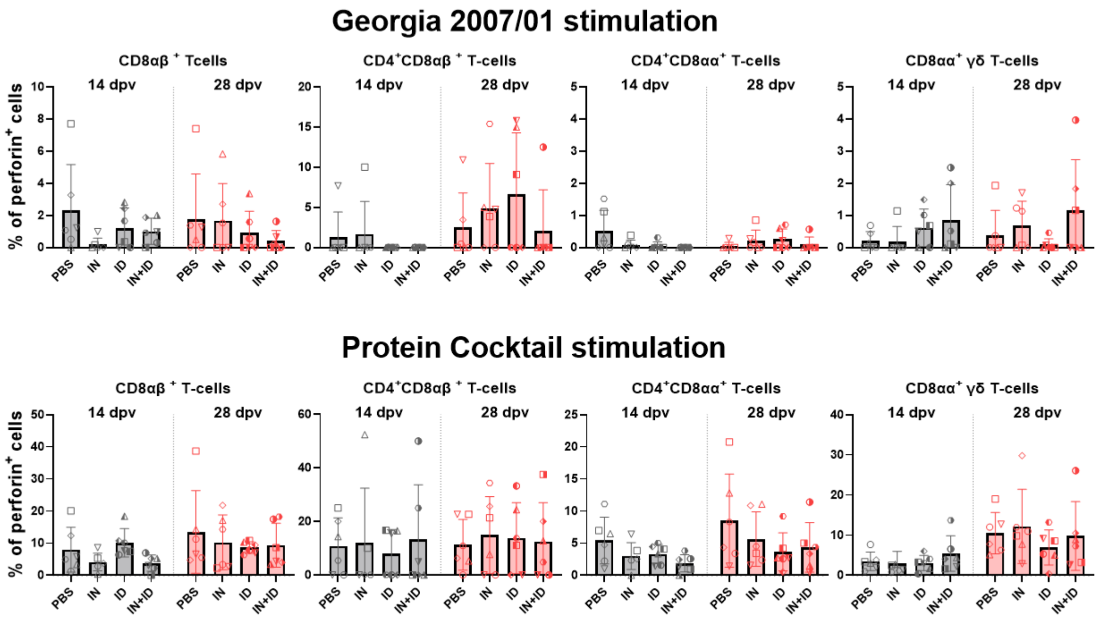


**Fig. S5.** Frequency of perforin-positive cytotoxic T-cell subsets following stimulation with Georgia 2007/1 or the vaccine cocktail. Percentage of perforin-positive T-cell populations among PBMCs collected at 14 and 28 dpv following *in vitro* stipulation with Georgia 2007/1 (A) or with the complete ASFV protein cocktail (B).

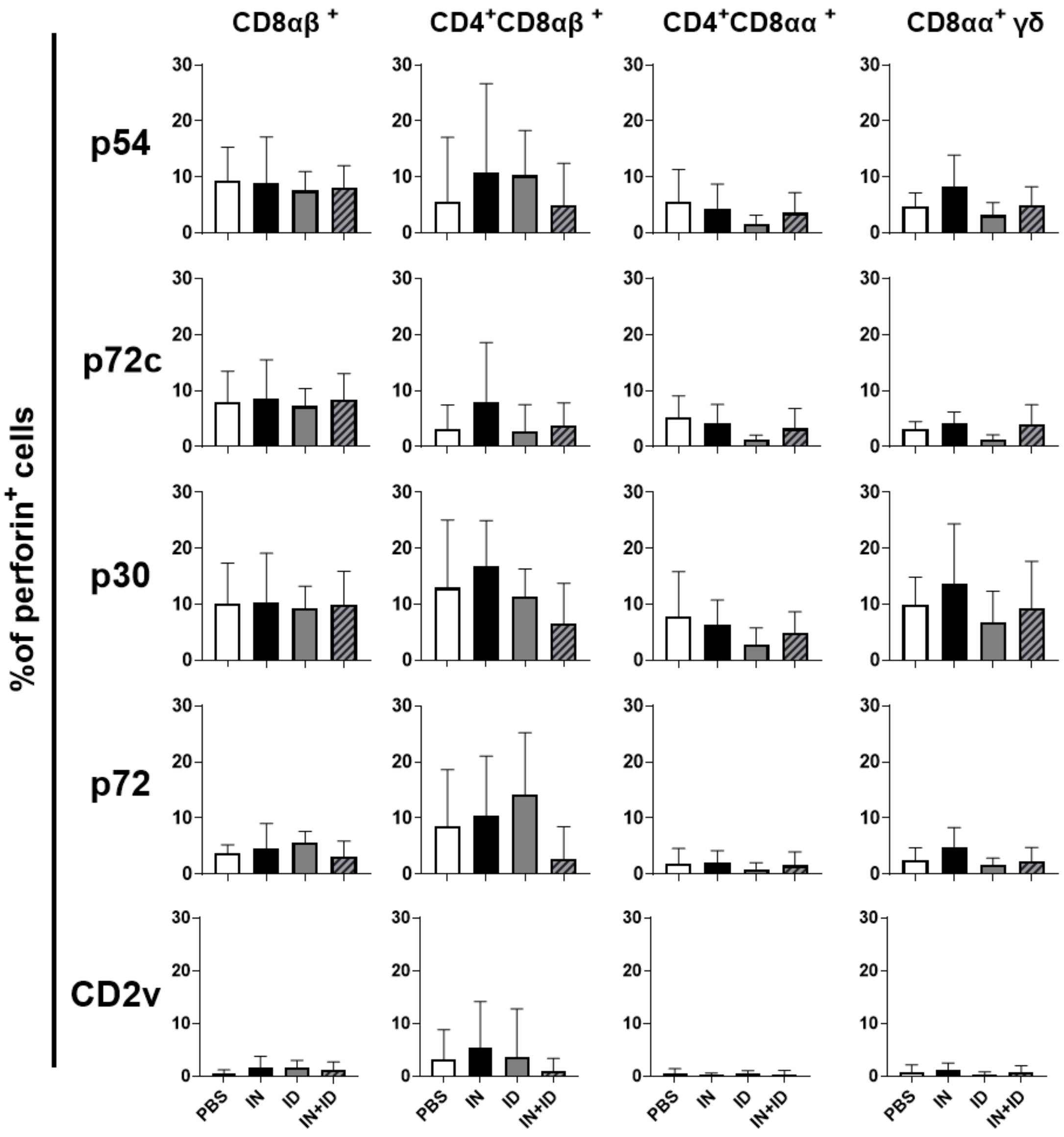


**Fig. S6.** Antigen-specific perforin-positive T-cell subsets. Percentage of perforin-positive T-cell populations at 28 dpv (0 dpc) following *in vitro* stimulation with the ASFV proteins included in the vaccine formulation.

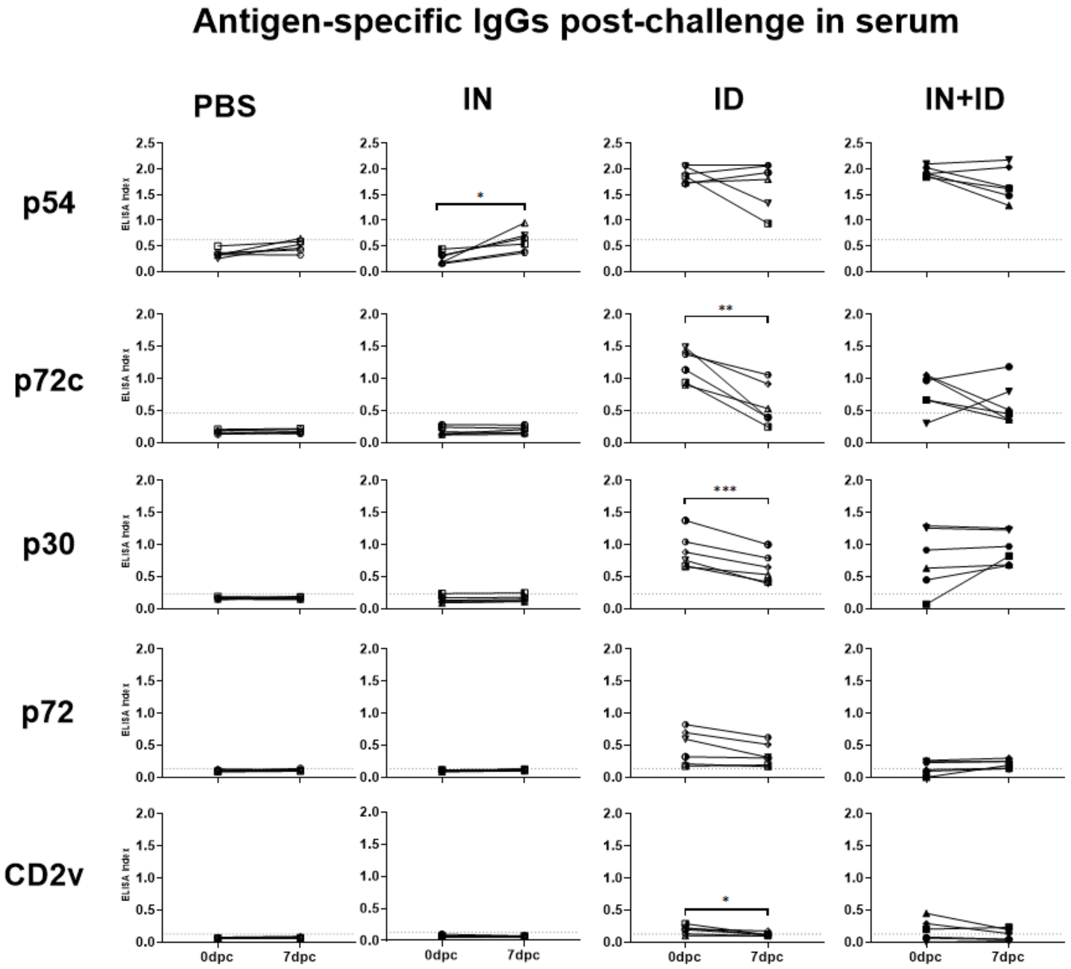


**Fig. S7.** Antigen-specific IgG responses following virulent ASFV challenge. Serum IgG antibodies against p54, p72c, p30, p72, and CD2v were measured by ELISA at 0 and 7 dpc in pigs vaccinated through the ID, IN and ID+IN routes. Statistical significance was determined by a paired t-test (*$P < 0.05$, **$P < 0.005$ and ***$P < 0.0005$).
